# Quantifying paddling kinematic through muscle activation and whole body coordination during maximal sprint of different duration on a kayak ergometer: a pilot study

**DOI:** 10.1101/2023.01.23.525293

**Authors:** Y.M. Garnier, P. M. Hilt, C. Sirandre, Y. Ballay, R. Lepers, C. Paizis

**Affiliations:** EA3920 Prognostic Factors and Regulatory Factors of Cardiac and Vascular Pathologies, University of Franche-Comté, 25000 Besançon, France; INSERM 1093-CAPS, Université Bourgogne Franche-Comté, UFR des Sciences du Sport, F-21000 Dijon and France; Centre for Performance Expertise, Faculté des sciences du sport, BP 27 877, Université de Bourgogne, Dijon F-21078 and France

**Keywords:** flatwater kayaking, body coordination, electromyography, pacing strategy

## Abstract

Paddling technique and stroke kinematics are important performance factors in flatwater sprint kayaking that require important energetic demand and high strength from the muscles of the trunk and upper limb. The various distance competed (from 200-m to 1000-m) requires the athletes to optimize their pacing strategy to maximize power output distribution throughout the race. This study aims to characterize paddling technique and stroke kinematics during two maximal sprints of different duration. Nine national-trained participants performed a 40-seconds and a 4-min sprint at maximal intensity on a kayak ergometer. The main findings demonstrated a significantly greater mean stroke power (237 ± 80 W vs 170 ± 48 W) and rate (131 ± 8 spm vs 109 ± 7 spm) during the 40-s sprint compared to the 4-min sprint. Athletes used an all-out strategy for the 40-sec exercise and a parabolic-shape strategy during the 4-min. Despite different strategies implemented and higher muscular activation during the 40-s sprint, no change in paddling technique and body coordination occurred during the sprints. The findings of the present study suggest that athletes constructed a well-defined profile that is not affected by fatigue despite a decrease in power output during the all-out strategy. Also, they regulate paddling kinematics during longer exercise with no change in paddling technique and body coordination.

## 1. Introduction

Flatwater sprint kayaking events are contested over distances from 200-m to 1000-m in single or crew boats composed of 2 or 4 athletes. In single kayaking, the world’s best race times ranged from 33-sec and 37-sec on 200-m until 200-sec and 230-sec on 1000-m for men and women, respectively (data from International Canoe Federation) which makes aerobic and anaerobic capacities as well as high upper body strength strong physiological factors of performance (Pickett et al. 2018; Bishop 2000; Forbes et al. 2009). Furthermore, the cyclic nature of the activity makes also the paddling technique an important performance factor that athletes should optimize to ensure the effectiveness of each blade stroke to propel the boat despite the persistent unbalance state resulting from the thinness of the boat.

From a global perspective, (Lopez Lopez et Ribas Serna 2011) identified a similar paddling technique among Olympic athletes who qualified for an “optimal stroke profile”. Qualitatively, (Limonta et al. 2010) reported longer stroke length developed by elite athletes compared to national-level athletes at the end of an incremental 4-min test. On 200-m sprint, (Pickett et al. 2021) pointed out that among the elite athletes, those of higher performance level also performed longer stroke length on 200-m races than elite athletes of a lower performance level. The strong negative relationship observed between stroke length and race time conducted by these authors highlights stroke length as a performance predictor of a 200-m sprint for elite athletes (Pickett et al. 2021). However, one should be cautious not to interpret these findings as increasing stroke length through a delayed exit of the blade from the water that would impair boat speed and consequently increase drag force (Gomes et al. 2015). Instead, international elite athletes of better performance level distinguishes from national level athletes by a greater range of motion in knee extension, increasing pelvis and trunk rotation which ultimately increased stroke length through a well forward catching phase (Limonta et al. 2010; Klitgaard et al. 2021). However, they maintain the exit phase once the blade reaches the vertical position to quickly initiate the contralateral catching phase while reducing ineffective portion of the stroke (Gomes et al. 2015). Particularly, (Gomes et al. 2015) reported that increasing stroke rate permitted a more rectangular force profile to be produced from the blade/water interaction, which results in a greater impulse during stroke and a higher mean boat velocity. Accordingly, the increase in mean stroke rate during 200-m races paralleled improvements in performances over the years (McDonnell, Hume, et Nolte 2013), which identifies stroke rate as a performance factor between international elite athletes and athletes of national level (Pickett et al. 2021).

Leg contribution in flatwater kayaking should also not be discarded. Providing the feet are strapped on the footrest, the kayakers performed push and pull actions with their legs to enhance pelvis and trunk rotation (Nilsson et Rosdahl 2016; Limonta et al. 2010; Klitgaard et al. 2021) (Nilsson et Rosdahl 2016) estimated to 0.2 seconds the duration of the synchronized period between leg extension and paddle thrust. Although of short duration, synchronization between leg extension, pelvis, and trunk rotation, as well as paddle thrust, needs to be finely refined to provide gain in boat velocity quantified from 6% (Begon, Colloud, et Sardain 2010, 201) to 16% (Nilsson et Rosdahl 2016).

It should, however, be mentioned that the vast majority of the studies describes paddling technique on a reduced number of stroke cycles and with no consideration about changes that could occur throughout a race. Only few studies analyze stroke kinematics over the time course of 200-m or 500-m maximal sprints (Pickett et al. 2021; Bertozzi et al. 2022; Vaquero-Cristóbal et al. 2013). Over short-distance sprint (i.e., 200-m), a reduction in boat speed and stroke rate was identified in young national-level paddlers (Vaquero-Cristóbal et al. 2013) and international and national level athletes (Pickett et al. 2021). (Vaquero-Cristóbal et al. 2013) also identified a reduction in stroke efficiency over the time-course of the 200-m sprint. On a 500-m sprint, (Bertozzi et al. 2022) also observed reductions in stroke length and velocity, while stroke duration remained constant. These elements shed light on the occurrence of fatigue that athletes face during a race to maximize their performance. Given the different distances competed in flatwater kayaking (i.e., from 200-m to 1000-m), athletes set up different pacing strategies to optimize power distribution throughout the sprint and limit the occurrence of fatigue (Jones et al. 2008; Abbiss et Laursen 2008). Therefore, pacing strategies represent a performance factor in flatwater sprint kayaking (Bishop, Bonetti, et Dawson 2002). These strategies also impact stroke peak power, power distribution or stroke rate, and physiological responses during sprint (Bishop, Bonetti, et Dawson 2002) (Therrien, Colloud, et Begon 2011). For instance, male athletes increased stroke rate and reduced stroke length to produce greater stroke impulse during paddling. Despite the lack of study regarding sprint of longer distance (e.g., 1000-m), one could suggest that the distribution of stroke rate or power during a 1000-m sprint race depends on the pacing strategy used. Consequently, athletes would also likely adapt their paddling technique (paddle displacement and body coordination) differently depending on the distance covered and their pacing strategy.

Adaptations in stroke kinematic and paddling technique can result from body co-ordination changes. (Bjerkefors et al. 2018) identified for instance that elite athletes increased shoulder flexion and adduction and performed trunk flexion and rotation of greater amplitude to increase power output during paddling. Also, (Bertozzi et al. 2022) evidenced that elite athletes increased lateral trunk flexion over a 500-m maximal sprint to compensate for the fatigue-induced reduction in shoulder range of movement amplitude. However, they did not detect a change in knee or ankle range of motion during the sprint (Bertozzi et al. 2022). In addition to motion capture, surface electromyography (sEMG) recordings can complete the investigation of body coordination related to paddling technique and evidence changes in the activation pattern of the muscles with the occurrence of fatigue. To date, the pattern of muscle recruitment during kayaking has, however been scarcely investigated, and limited to comparison in shoulder’s muscles recruitment between on-water kayaking and ergometer (Trevithick et al. 2007) or the lower limb in with-water races (Murtagh et al. 2016). In this context, it remains currently unknown how athletes adapt their paddling technique during a prolonged sprint when facing fatigue.

This study aimed therefore to identify whether national-level athletes adapted different paddling technique and stroke kinematics during simulated maximal sprint of different durations. Additionally, we intended to identified how the different pacing strategy employed by athletes impacts the occurrence of fatigue and its effects on body coordination and muscle activation patterns. We hypothesized that athletes would use a more conservative pacing strategy during the 4-min sprint than an all-out strategy employed during the 40-sec sprint, limiting kinematic and muscle coordination changes.

## 2. Materials and Methods

### 2.1. Participants

Nine experienced young kayak athletes (2 females, age: 18 ± 3 years; BMI: 22.2 ± 2.0 Kg.m-1; training experience: 8.0 ± 5.2 years) participated in this study. All the athletes competed at least at national level, and two were members of the U18 French kayak team. The study conformed to the standard set by the World Medical Association Declaration of Helsinki “Ethical Principles for Medical Research Involving Human Subjects” (2008).

### 2.2. Experimental design

Participants visited the laboratory in a single session to performed two maximal efforts on a flywheel kayak ergometer (Dansprint PRO kayak ergometer, Dansprint ApS, Denmark). All participants were experienced with paddling on an ergometer for their personal training. The distance between the seat and the foot bar was set for each athlete to correspond with the settings of their kayak. The flywheel resistance was adjusted with account of the athlete’s weight to reproduce on-water drag forces following the manufacturer’s prescription (http://www.dansprint.com). Participants were instructed not to undertake vigorous training 24h before the experimentation.

Before the experimental acquisitions, participants performed a warm-up including 6-min paddling at an intensity of 70% of their maximal perceived effort, 2-min at 90% of their maximal perceived effort, and three 10-s maximal efforts. Recovery periods of 2-min followed the 6-min and the 2-min sets, and 1-min rest between the 10-s efforts. Then, participants performed two maximal exercises of different duration 40-s and 4-min to correspond with average duration for 200-m and 1000-m, with 15-min of recovery in-between. Athletes were asked to perform the highest distance they could during each bout, with no instruction about pacing strategy and no feedback on their performance.

### 2.3. Stroke kinematics

Stroke rate and power were recorded stroke-by-stroke and stored for off-line analysis using the ergometer software (Dansprint analyser V1.12). Data were averaged over 15 complete cycles defined by the position of the paddle tip (McDonnell, Hume, et Nolte 2012) and analyzed at three-time points corresponding to the beginning (T1), the middle (T2), and the end (T3) of each exercise.

### 2.4. Surface Electromyography (sEMG)

EMG activity was recorded during exercises using wireless EMG probes (Cometa systems, Italy) at a frequency of 1000 Hz. Pairs of pre-gelled surface electrodes (10 mm diameter) were applied to the midpoint of the palpated muscle belly along the muscle fibers with a 2 cm centre-to-centre interval. Ten upper and lower limb muscles were investigated: anterior, middle, and posterior deltoids (Del), upper trapezius (Trap), pectoralis major (Pec), latissimus dorsi (Lat), biceps brachii (Bic), triceps brachii (Tri), rectus abdominis (Rect) and vastus lateralis (Vast). The activity of the three portions of the deltoids was average to reflect the activity of the global deltoids. EMG activity was averaged over 1 stroke cycle and analyzed with automatic processing (Matlab, Mathworks, USA).

### 2.5. Motion capture

Three-dimensional markers trajectory was recorded using seven infra-red cameras sampling at 200 Hz (Vicon T-series, Vicon motion systems Ltd. UK). Nineteen retro-reflective markers (diameter = 20mm) were applied on the body to identify the trunk, upper- (forearm, arm, shoulder girdle) and lower-limb (pelvis, thigh, leg) and the paddle (see Fig. 1). Based on the 3D movement of these markers, we defined an elevation angle for each segment, as the angles between each segment and the vertical axis (defined by gravity-extrinsic reference). In addition, the amplitude of the paddle displacement in the vertical and horizontal plans (X, Z) was also evaluated. To assess the coordination between the different segments during the movement, a PCA was performed (for more details on the computation, see (Berret et al. 2009; Paizis et al. 2008) upon the time series of the 10 elevation angles (upper-arm, arm, trunk, leg, lower-leg for each hemibody). Here we used, as coordination value, the percentage associated with the three first principal components for 15 double-stroke cycles at the beginning, the middle, and the end of each test (40sec vs 4min).

**Figure 1.**
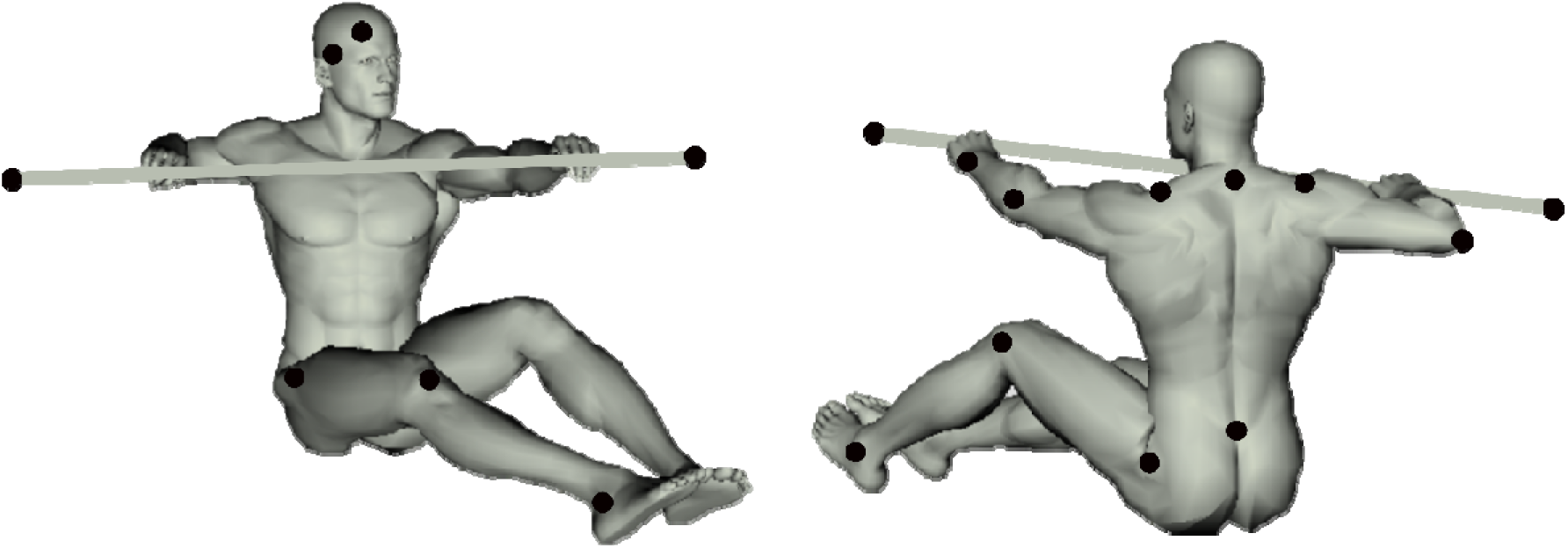
Placement of the retro-reflective markers on the participants.

### 2.6. Statistical analysis

All data are presented as mean ± standard deviation (SD) in text, figures and tables. The nature of the distribution was assessed for all variables using the Shapiro-Wilk test to ensure proper use of parametric ANOVA. A Greenhouse-Geisser correction to the degree of freedom was applied when the sphericity of the data was violated. A two-way 2 × 3 ANOVA was used to test the effect of condition (40-sec vs. 4-min exercise) and time (T1 vs. T2 vs. T3) on EMG activity, kinematic and stroke rate and power. When significant, main effect and condition × time interaction were followed-up with Tuckey HSD test. Effects sizes are reported as partial eta squared (ηp^2^) and Cohen’s dz, the latter being calculated from the mean and standard deviation of the variables, and the correlation between these variables using G*Power (version 3.1, Universität, Düsseldorf, Germany). Statistical analysis was performed with Statistica (StatSoft France, version 7.1, STATISTICA). The significance level was set at 0.05 (two-tailed) for all analysis.

## 3. Results

### 3.1. Stroke rate and power

The significant condition × time interaction for the stroke rate (*p = 0.007; ηp^2^ = 0.464*) revealed significantly lower stroke rate during the 4-min exercise compared to the 40-s at each time point (*all p < 0.001; all dz > 0.921;* ***see fig. 2A***). Stroke rate was significantly lower at T2 compared to T1 and T3 during the 4-min exercise (*all p < 0.018; all dz > 1.528*), with no difference between T1 and T3. Stroke rate remained constant during the 40-s exercise (*all p > 0.747; all dz < 0.680*). The ANOVA also detected a condition × time interaction on stroke power (*p < 0.001; ηp^2^ = 0.598;* ***see fig 2.B***). Power was greater during the 40-s exercise compared to the 4-min at each time point (*all p < 0.013; all dz > 0.508*) and significantly higher at T1 than T2 and T3 in each condition (*all p < 0.031; all dz > 1.148*). Power increased from T2 to T3 during the 4-min exercise (*p = 0.005; dz = 1.521*), while remaining constant for the 40-s exercise (*p = 0.971; dz = 0.265*).

**Figure 2.**
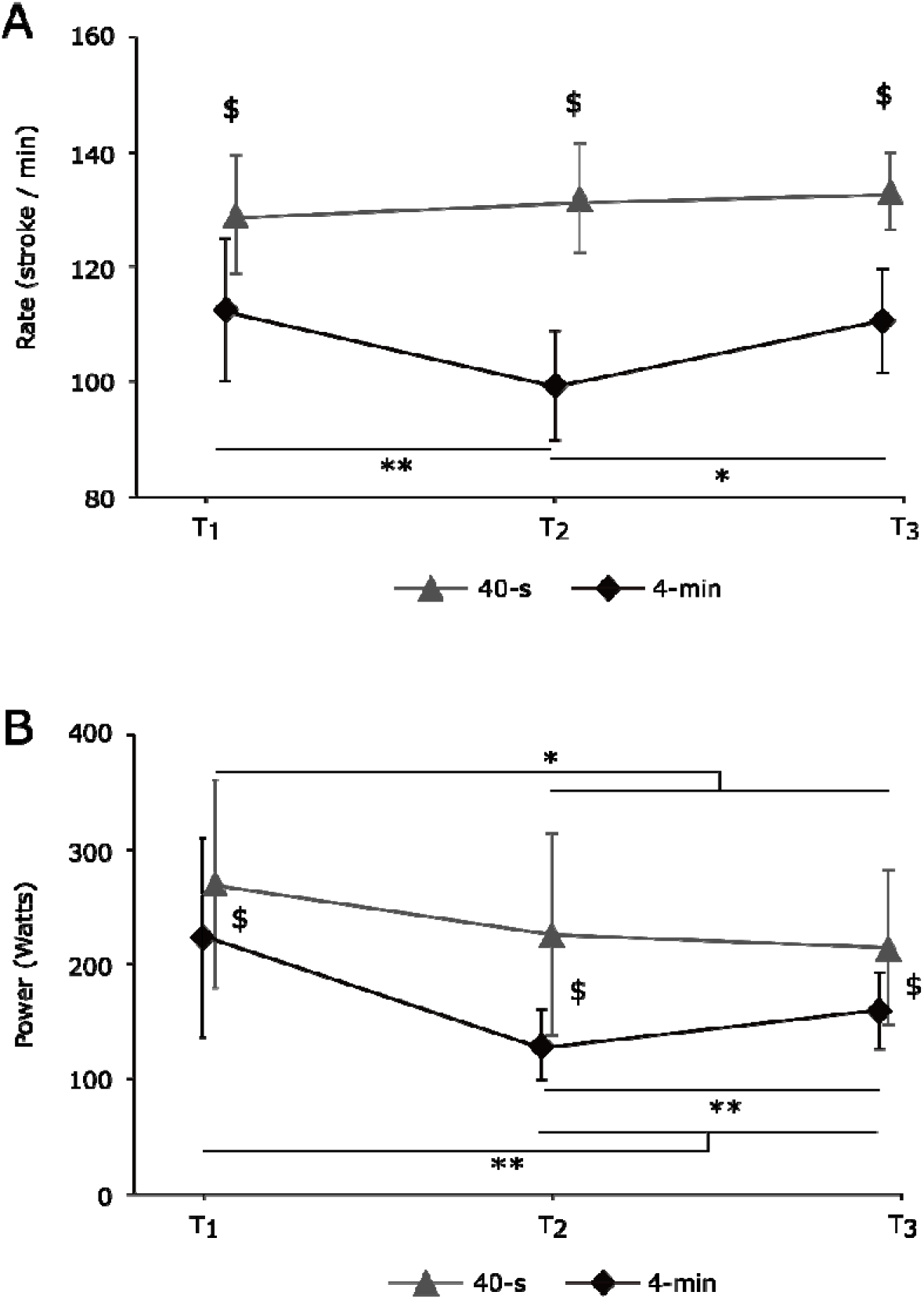
Stroke rate (A) and power (B) recorded during the 40-s and 4-min sprints (N = 9; mean ± SD).

### 3.2. Electromyographic activity

**Figure 3**. presents EMG activity of each muscle at the different time points during the 40-s and the 4-min exercise. Due to a technical issue during recording, EMG data of one participant has been removed from analyses, and only data from 8 participants have been considered. A significant main effect of time was detected for *Pec, Trap*, and *Lat* muscles (*all p < 0.006; all ηp^2^ > 0.516*), with no condition or condition × time interaction (*all p > 0.057; all ηp^2^ < 0.434*). All three muscles demonstrated greater EMG activities at T2 compared to T3 (*all p < 0.004; all dz > 1.336*). EMG activity was also greater at T2 compared to T1 for the *Trap* muscle (*p < 0.001; dz = 1.675*).

A significant condition × time interaction was detected for *Bic, Tri, Del, Rect* and *Vast* muscles (*all p < 0.049; all ηp^2^ > 0.350*). The Rect muscle showed a significantly greater EMG activity at T1 and T3 during the 40-s than during the 4-min exercise (*all p < 0.026; all dz > 0.809*). All other muscles (i.e. *Bic, Tri, Del*, and *Vast*), showed a greater EMG activity during the 40-s than the 4-min exercise at each time points (*all p < 0.001; all dz > 0.873*). The Vast muscle showed no change of the EMG activity over time during both exercises (*all p > 0.062; all dz < 0.917*). During the 4-min exercise, the Tri and Del muscles showed greater EMG activities at T2 compared to T1 and T3 (*all p < 0.018; all dz > 1.218*), and EMG activity of the Bic was greater at T2 compared to T3 (*p = 0.021; dz > 0.764*). No other change was observed.

**Figure 3.**
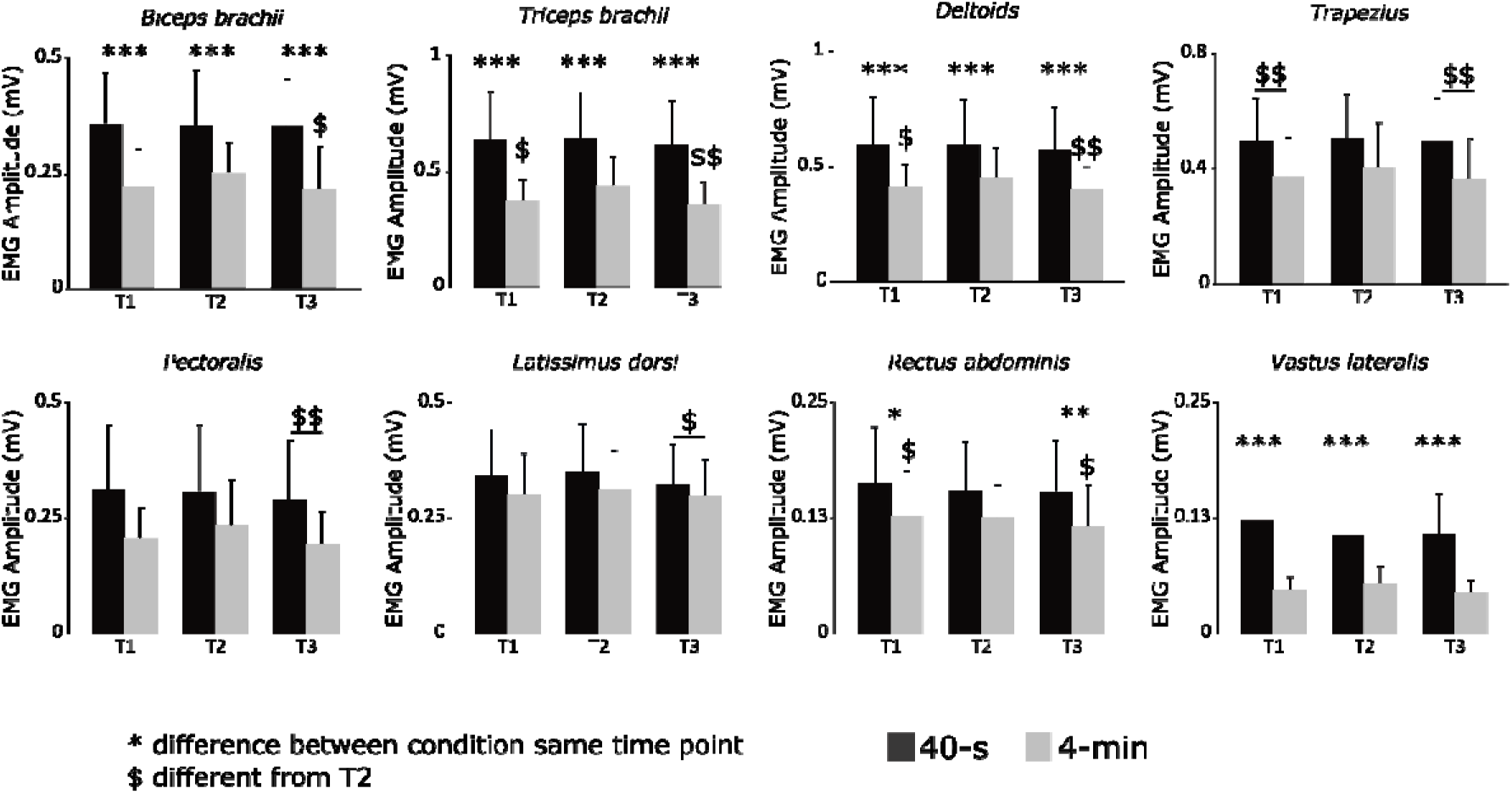
EMG amplitude of muscle activations recorded at the beginning (T1), middle (T2) and at the end (T3) of each sprint (N = 8; mean ± SD).

### 3.3. Kinematic data

***Table 1*** showed the mean elevation angle of the upper arm, arm, and trunk calculated for the two conditions. Due to technical problems with reflective markers, only complete data from 4 subjects have been included in kinematic analysis. The ANOVA revealed neither main effect of the condition or time (*all p > 0.271; all ηp^2^ < 0.353*), nor interaction (*p = 0.279; ηp^2^ = 0.364*) for elevation angles coordination.

**Table 1.** Mean and standard deviation, across subjects, of the main kinematic parameters for the 40 sec and 4 min condition.

|  | 40 sec. |  | 4 min. |  |
| --- | --- | --- | --- | --- |
|  | start | end | start | end |
| **Right Hemibody** |  |  |  |  |
| Elev. Angle upper arm (deg.) | 67 ± 2 | 64 ± 2 | 67 ± 3 | 65 ± 2 |
| Elev. Angle arm (deg.) | 70 ± 4 | 72 ± 3 | 68 ± 3 | 68 ± 2 |
| Elev. Angle trunk (deg.) | 14 ± 2 | 14 ± 1 | 14 ± 1 | 14 ± 1 |
| Amplitude Paddle Vert. (cm) | 164 ± 15 | 167 ± 15 | 160 ± 14 | 162 ± 7 |
| Amplitude Paddle Hrzt. (cm) | 61 ± 9 | 63 ± 8 | 64 ± 11 | 66 ± 12 |
| **Left Hemibody** |  |  |  |  |
| Elev. Angle upper arm (deg.) | 67 ± 2 | 64 ± 2 | 67 ± 3 | 65 ± 2 |
| Elev. Angle arm (deg.) | 70 ± 4 | 72 ± 3 | 68 ± 3 | 68 ± 2 |
| Elev. Angle trunk (deg.) | 14 ± 2 | 14 ± 1 | 14 ± 1 | 14 ± 1 |
| Amplitude Paddle Vert. (cm) | 158 ± 2 | 154 ± 3 | 158 ± 2 | 153 ± 7 |
| Amplitude Paddle Hrzt. (cm) | 67 ± 12 | 71 ± 11 | 74 ± 7 | 74 ± 5 |
| Coordination (%) | 94 ± 2 | 94 ± 2 | 94 ± 2 | 93 ± 3 |

## 4. Discussion

This study aims at describing kinematic and muscle activity of the paddling technique in young, experienced paddlers during maximal sprints of different duration on a kayak ergometer, and to analyze how athletes adapt their pacing strategy depending on race duration. In line with our hypothesis, the results from stroke power highlighted that the athletes use a more conservative pacing strategy during the 4-min sprint than an all-out strategy employed during the 40-sec sprint, with higher muscular activity in biceps brachii, triceps, deltoids, rectus femoris, and vastus lateralis muscles during the 40-sec sprint. Despite these differences not found differences in kinematic and muscle coordination.

Mean power (237 ± 80 W and 170 ± 48 W) and stroke rate (131 ± 8 spm and 109 ± 7 spm) recorded during the 40-s and 4-min sprint, respectively, are similar to that reported for national-level athletes by (Pickett et al. 2021) on 200-m and somewhat lower than that reported for international athletes by (Bjerkefors et al. 2018). Maximal and mean power and stroke rate were significantly lower during the 4-min bout compared to the 40-s sprint despite the same resistance imposed by the ergometer. During the 40-s sprint, stroke rate remained constant while stroke power decreased immediately after the start (i.e. from T1 to T2 and T3), with no further decrease between T2 and T3. During the 4-min sprint, stroke rate and power were higher at T1 compared to T2 and T3, and both parameters increased from T2 to T3. These elements showed that athletes implemented two different pacing strategies during exercise. Athletes implemented an all-out strategy during the 40-s sprint characterized by a greater power developed at the start and then maintained as long as possible, while an even-pace strategy characterized by a parabolic-shape profile was implemented during the 4-min exercise (Bishop, Bonetti, et Dawson 2002; Abbiss et Laursen 2008). These two strategies previously reported for well-trained kayakers (Bishop, Bonetti, et Dawson 2002; Pickett et al. 2021) and rowers (Dimakopoulou et al. 2018), would improve performance through optimization of the balance between power production and occurrence of fatigue. Specifically, an all-out strategy would enable athletes to achieve the greater amount of work they could above the critical power before a high level of fatigue limiting their performance (Jones et al. 2008). On the contrary, athletes would implement an even-pace strategy based on their own experience to limit disturbance in muscle intracellular milieu (Bishop, Bonetti, et Dawson 2002), delaying therefore the occurrence of fatigue (Dimakopoulou et al. 2018) allowing them to limit decrease in boat speed or even increase it.

Performance in sprint flatwater kayaking also stands upon the acquisition of a fine paddling technique that ensures the effectiveness of the paddling stroke throughout the race. In line with the “optimal stroke profile” identified by (Lopez Lopez et Ribas Serna 2011) that international elite athletes share, present findings showed the same paddling profile implemented by the athletes during both sprints (see ***fig. 4***). Furthermore, no change in this pattern was evidenced during the race despite different pacing strategies. Previously, well-trained and elite athletes also demonstrated synchronized actions between the upper and the lower limb (Bjerkefors et al. 2018; Nilsson et Rosdahl 2016). Our findings showed that elevation angles and body coordination were similar between the 40-s and the 4-min sprint and were similar between left and right sides of the body as ever reported by (Bjerkefors et al. 2018). Hence in accordance with (Therrien, Colloud, et Begon 2011), the present findings suggest that well-trained athletes kept the same paddling technique despite different stroke rate between sprints.

**Figure 4.**
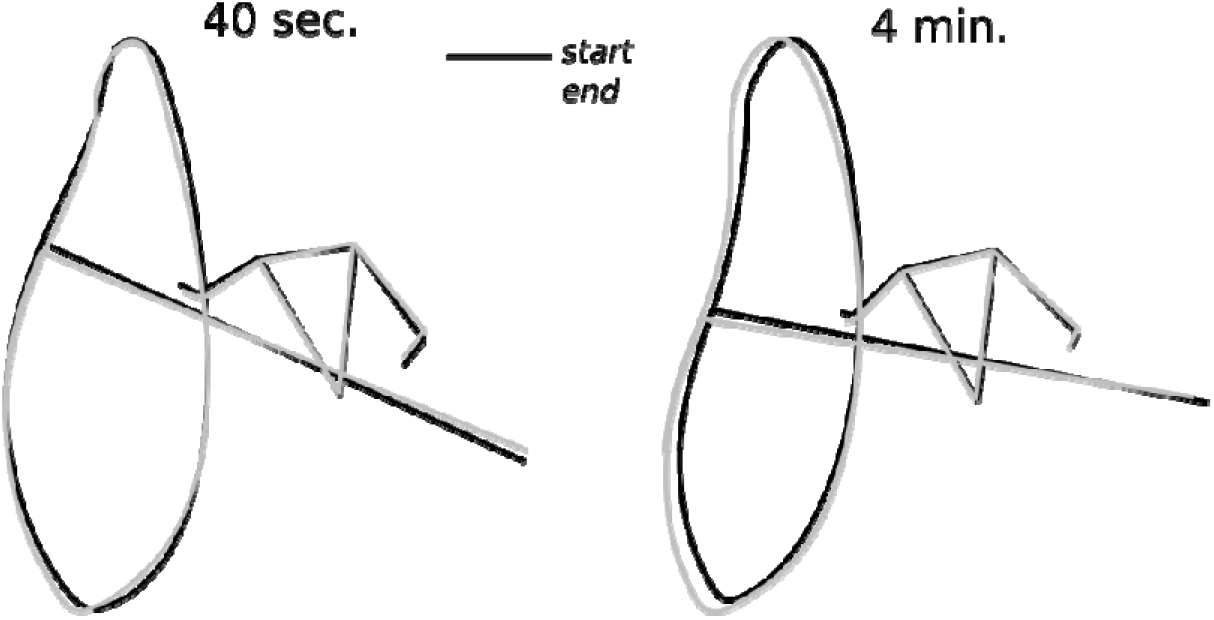
Mean start posture and paddle tip trajectory for the 15 first cycles (black) and 15 last cycles (grey) of the 40 sec (left) and 4 min (right) condition, for a typical subject.

Surface EMG was also recorded in our study to characterize the impact of different pacing strategies and the occurrence of fatigue on muscle activation patterns. Muscle activities of the upper- (i.e. Bic, Tri, Del) and lower limbs (i.e. Vast) were consistently h gher during the 40-s compared to the 4-min sprint. The greater mean activity of these muscles during the 40-s corroborates the higher power output recorded during sprint. Specifically, the increase in mean EMG activity of the biceps brachii would transcribe a greater force level developed to resist against elbow extension during the pulling phase (biceps brachii). In contrast, increase in triceps brachii activity would limits elbow flexion of the aerial limbs to ensure the effectiveness of the opposite pulling phase (Fleming, Donne, et Fletcher 2012; Bjerkefors et al. 2018). The greater activity of the vastus lateralis muscle recorded during the 40-s reflects the prominent role the lower limbs play in force transmission in flatwater kayaking (Nilsson et Rosdahl 2016; Begon, Colloud, et Sardain 2010). However, most of the investigated trunk muscles (i.e. Trap, Pec, Lat) did not demonstrate different EMG activity between sprint modalities, excepted a greater activity for the rectus abdominis at T1 and T3 during the 40-s compared to the 4-min. The trapezius and the latissimus dorsi muscles control for shoulder extension and internal rotation during kayak paddling (Trevithick et al. 2007). The similar magnitude of EMG activity between sprint modalities for these two muscles suggest that the greater power output produced during the 40-s did not increase shoulder instability. Together, these findings suggest that athletes adapt to the greater power output mainly by increasing muscular force to ensure the effectiveness of their paddling technique rather than change in paddling tip kinematic (e.g. pulling stroke length).

Flatwater race sprint kayaking performance is a complex process depending on numerous factors such as athletes’ level, inter-individuals’ history, balance and distance. Additionally, it would be of interest to explore in real conditions the muscular activations associated with the kinematics and body coordination with a larger sample size in different distances in a future study in order to optimize performance during single boat race and ensure optimal stroke synchronization in crew boat.

## 5. Conclusion

The main findings of this study showed that athletes used different pacing strategy between a 40-s and a 4-min maximal sprint. Specifically, athletes implemented an all-out strategy during the 40-s sprint in order to achieve the greater work above the critical power before the occurrence of neuromuscular fatigue, while employing an even-paced strategy to avoid excessive accumulation of fatigue early during the race. In spite of these two marked different strategies, athletes kept a similar paddling technique that could be defined as the “optimal stroke profile” ever reported in previous studies. Only increase in mean activity of upper- and lower limbs muscles was detected when comparing the 40-s to the 4-min sprint, which could be related to an increase in force to resist against the greater stroke power and maintain the effectiveness of their paddling technique. Therefore, we encourage practitioners to develop the most suitable strategy relatively to their distance race and give more attention in strength training of biceps brachii, triceps brachii, deltoids, rectus femoris and vastus lateralis which were more activated to produce more power for a similar paddling technique and body coordination.

